# Postglacial Genetic Legacies, Demography, and Climate Change: Setting Conservation Priorities for Silver Fir

**DOI:** 10.1101/2024.10.07.616973

**Authors:** Francisco Balao, Marc Ríos-Cadenas, José Manuel Sánchez-Robles, María Teresa Lorenzo, Juan C. Linares, Anass Terrab

## Abstract

**Aim:** Climate change challenges the adaptive capacity of several tree species. This study explores the genetic diversity and past range dynamics of silver fir (*Abies alba*) with the objective of set hotspots of population diversity and priorities for conservation based on climate change-induced dieback.

**Location:** Mediterranean, Europe

**Methods:** We perform genome-wide analysis by RAD-seq to range-wide 26 *A. alba* populations.

**Results:** We identified two lineages in the southern range, formed by populations from the Apennine chain, from the west side, and those from the Balkans and Carpathians, from the east side. Northern range populations showed two lineages, with definite genetic differences between the westernmost populations from those from the eastern. Populations from the Northern Balkans and the NE Apennine were situated in the intermediate positions. Regarding demographics, southern populations remained stable as glacial refuges, while northern populations experienced a population decline during the middle Pleistocene due to glaciations. Although the demographic events that shaped the current spatial structure of genetic diversity of silver fir go back to the Pleistocene, human pressure during the Holocene likely led to an abrupt range decline, while recent drought events have caused widespread dieback.

**Main Conclusions:** We provide a generic framework for setting silver fir conservation priorities based on the phylogeographical lineages and current dieback symptoms induced by climate change. Our results support that rear edge populations might be disproportionately important for the adaptive capacity of silver fir under a climate change scenario.

## 1 INTRODUCTION

While climate change constrains the persistence of numerous tree species (Allen et al., 2015; McDowell et al., 2020), the biogeography and genetic diversity of populations could be key to achieve local adaptations (Aitken et al., 2008). As regards biogeography, the response of species to changing environments is likely to be determined by population responses at range margins, where rear edge populations may be a long-term store of genetic diversity (Davis & Shaw, 2001; Hampe & Petit, 2005). Further, the relationship between standing genetic diversity and adaptive phenotypic characteristics may be essential to cope with climate change (Alberto et al., 2013; Barrett & Schluter, 2008).

Range-wide patterns of population genetic diversity are related to past climate-driven range dynamics. In the Northern Hemisphere, this standing genetic diversity was strongly influenced by post-glacial gene flow (Davis & Shaw, 2001; Hewitt, 2000, 2004). Hence, tree populations have been subject to cyclical patterns of extinction and dispersal, driven by the dynamics of glacial periods. During glaciations, forest species contracted their ranges, taking refuge in southern disjunct refugia, and subsequently extended their range northwards in response to post-glacial climatic amelioration (Hewitt, 1999). The main Mediterranean peninsulas would have acted as full-glacial refugia for temperate tree populations but northern have been suggested for some boreal and nemoral European tree species (Medail & Diadema, 2009; Svenning et al., 2008; Tzedakis et al., 2002). These refugia have significantly influenced the genetic structure and diversity of areas recolonized after the glaciations (Hewitt, 2000). Precise identification of glacial refugia is essential for conservation strategies, as these locations might retain the core of species’ genetic diversity (Fady et al., 2022; Hewitt, 2004; Keppel et al., 2012). Furthermore, these areas are crucial for the long-term preservation of biodiversity and face threats from rapid environmental changes in the Mediterranean region (Sánchez-Salguero et al., 2017). However, the precise identification of these refugia, and consequently the delineation of postglacial migration routes, remains elusive for certain species due to inherent methodological challenges (Willis & Van Andel, 2004).

Firs (*Abies*, Pinaceae) are among the dominant trees of the temperate-cool forests of the Northern Hemisphere, including North America, East Asia, and the Mediterranean Basin (Farjon & Rushforth, 1989). A recent study focusing on the evolutionary history of the Circum-Mediterranean firs (CMF) by Balao et al. (2020) supports the idea that all Mediterranean firs form a single, monophyletic group. Furthermore, this research evidences two lineages within the CMF, which align with the sections previously identified as *Abies* and *Piceaster* (Farjon & Rushforth, 1989). Additionally, this study suggested an early diversification of the Mediterranean firs in the late Oligocene-Early Miocene coupled with both ancient and recent admixture resulting from secondary contacts. This underscores the crucial role of historical climatic events in the speciation process and genomic architecture of the species in this fir lineage. Among the CMF, silver fir (*Abies alba* Mill.) is the only species with a widespread range, while the remaining taxa are mostly allopatric, with the distributions often confined to very limited areas (Linares, 2011).

The current understanding of the glacial and postglacial history of *A. alba* is based on paleoecological data and extensive genetic studies conducted only at regional level or using a limited number of molecular markers (e.g. Fady et al., 1999; Gömöry et al., 2012; Hussendörfer, 1999; Kempf et al., 2020; Longauer, 2001, 2003; Piotti et al., 2017; Teodosiu et al., 2019; Vendramin et al., 1999; among others). During the Last Glacial Maximum, *A. alba* is hypothesized to have found refuge primarily in two principal locations, the northern Apennines, and the southern Balkan Peninsula (Figure 1). Further, refugia in the southern Apennines and the Pyrenees has been proposed (Liepelt et al., 2002, 2009; Litkowiec et al., 2016; Terhürne-Berson et al., 2004), although these populations might have remained largely isolated, resulting in only limited expansions and, consequently, they would be genetically distinct (Fady et al., 1999). Nevertheless, the Pyrenees refugium has been considered uncertain by other authors (Reille & Lowe, 1993). Additionally, palaeogeological evidence also suggests different ice-age refugia of silver fir in two majors glacial refugia located in northern Apennines and northwestern Greece (Cheddadi et al., 2014).

**FIGURE 1.**
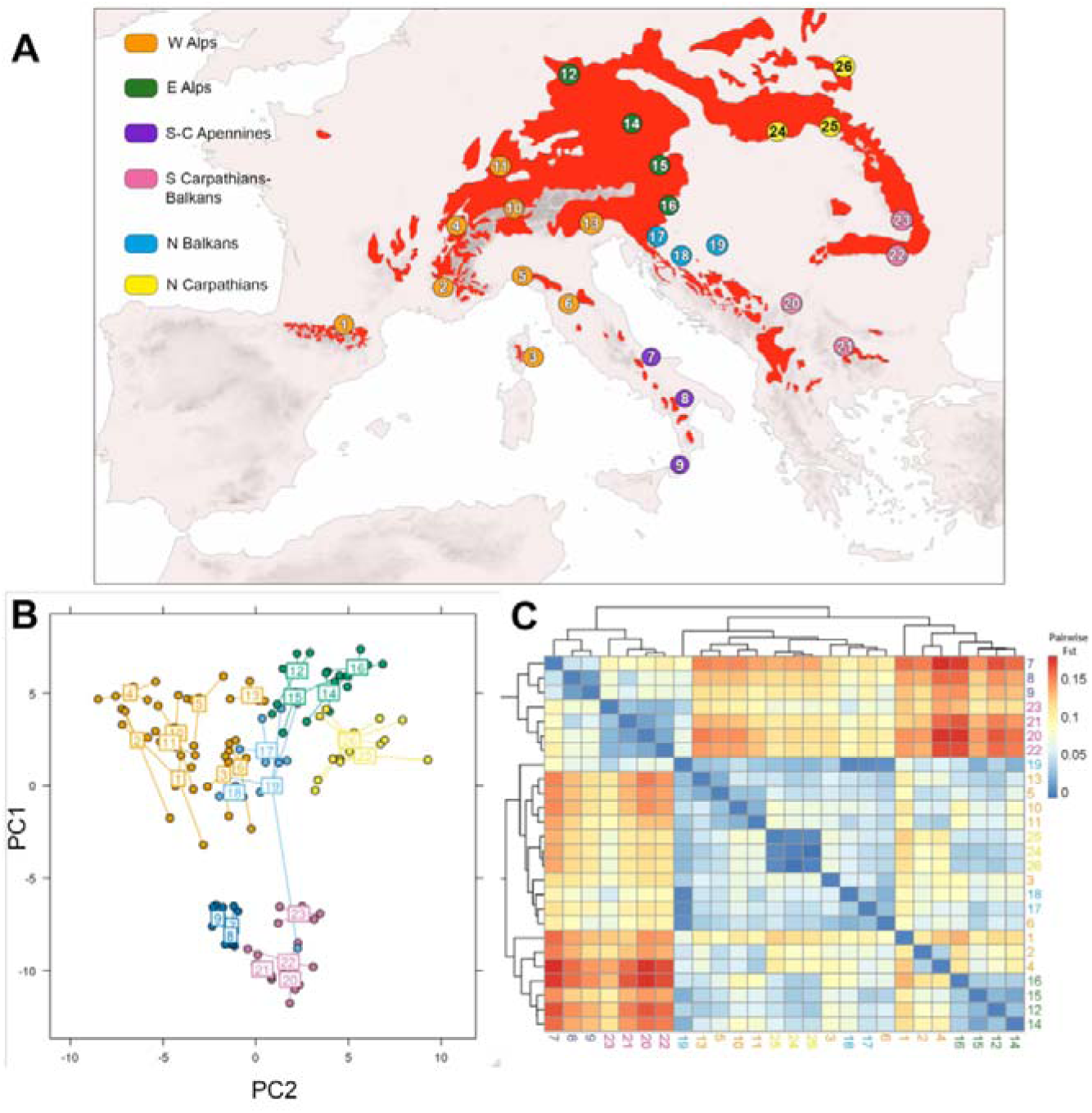
A) Map showing sampled populations within the distribution range of *Abies alba*, marked in red (Source: EUFORGEN). Numbers indicate the investigated population locations (detailed in Table 1), with the main geographic regions differentiated by color. B) PCA reveals regional segregation across 26 populations along north-south and west-east axes. C) Heatmap displays *F*_ST_ genetic distances, warmer colors signal higher distances, cooler colors lower.

**TABLE 1.**
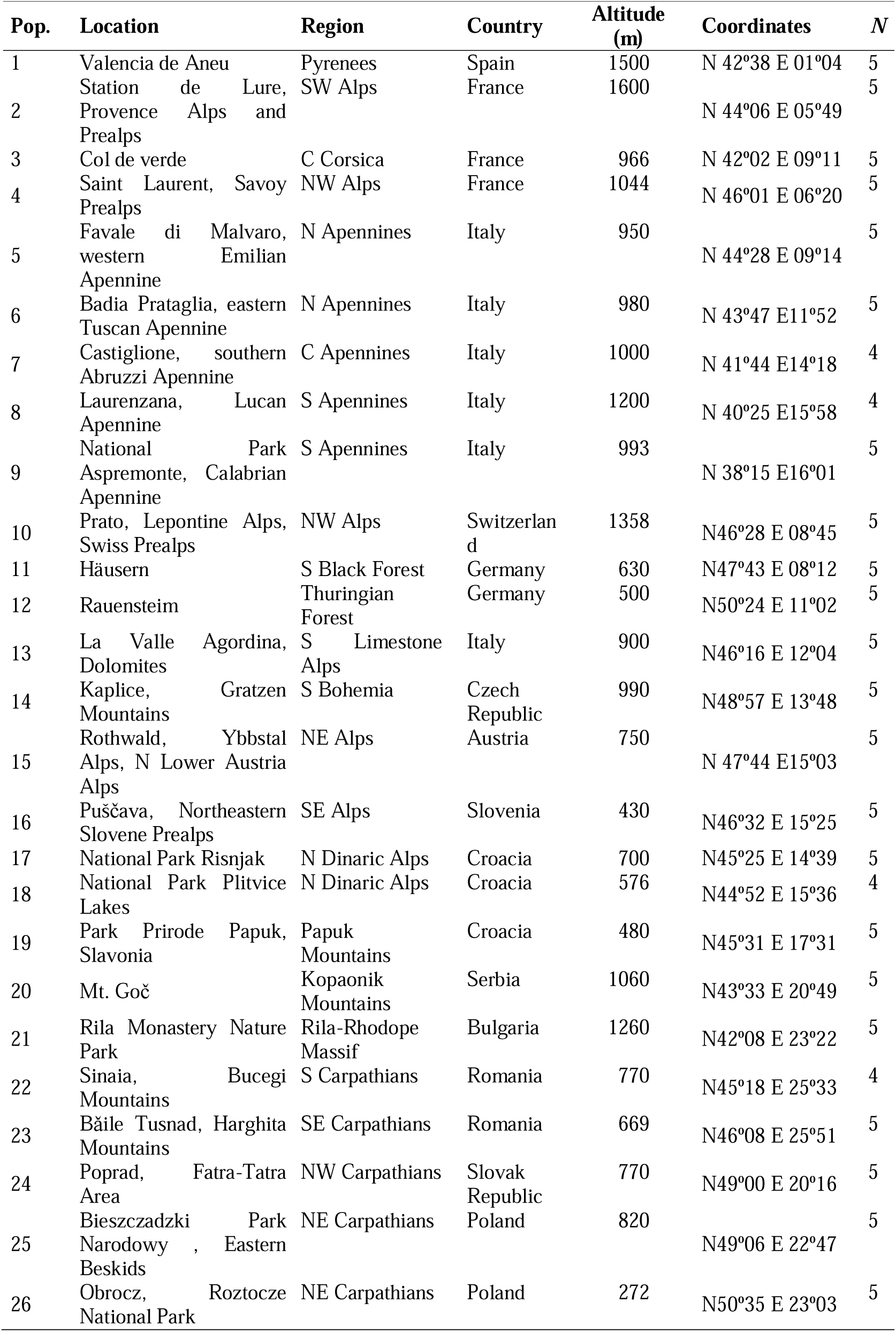
The 26 *Abies alba* populations sampled. *N* represents the number of individuals analysed.

The potential pathways for postglacial recolonization and zones of introgression for *A. alba* have been also under debate (Konnert & Bergmann, 1995; Liepelt et al., 2002). It is proposed that the recolonization of northwestern, central, and northeastern European regions occurred via two primary postglacial routes: 1) originating from the central Apennines, and 2) emanating from the southern Balkan Peninsula (Figure 1). The northwards colonization from N Apennines would happen via two subroutes as suggested by Karl (1980). In the “West Alpine route”, silver fir move along the Ligurian mountains into the southeastern French Alps and then colonized the Jura, Vosges mountains and Black Forest. In the “East Alpine route”, outflanking the Alps in the east and led to the Bavarian, Bohemian and Thuringian Forests. The second major colonization route linked south Balkan Peninsula silver fir populations to northern areas via two additional subroutes. A “West Balkan route” would move along the Dinaric Alps towards the eastern Alps. In the “East Balkan route” colonizing the Carpathians via S Bulgaria. Despite most studies outlined the importance of range shifts (Williams et al., 2004), the potential for genetic responses has often been neglected. Hence, to date, no study has thoroughly investigated the genomic and demographic consequences of glacial cycles throughout the full distribution range of silver fir, to validate these proposed refugia and post-glacial migration routes.

In this study, we harness the analytical power of restriction site-associated DNA sequencing (RAD-seq) to perform an exhaustive genome-wide analysis across 26 silver fir populations, each carefully selected to represent the entire current distribution range. Our approach is fundamentally aimed at testing key hypotheses concerning the historical number of refugia and the principal postglacial colonization pathways that have shaped the present-day distribution of *A. alba*. We specifically aimed to 1) delineate the phylogenomic relationships among populations of silver fir, 2) identify and characterize genetic discontinuities and contact zones between distinct genetic lineages, thereby elucidating the landscape genetic structure, and 3) discuss conservation priorities from the inference of past demography and current vulnerability to climate change.

## 2 MATERIALS AND METHODS

### 2.1 Sample collection and DNA extraction

*A. alba* currently inhabits in Central Europe, on the Suisse plateau and in South and Eastern Germany as well as in the Czech Republic and Austria. There are patent populations in the Pyrenees, Southern Alps of Northern Italy and the Eastern Alps, the Carpathians and Albania. It is also found more sporadically in Eastern France, on the Massif Central, and in the Apennines. Stands of silver fir are present in the Dinaric Alps which are continuously connected towards the Rhodope Mountains in Bulgaria and Greece (EUFORGEN, 2011). We sampled 26 autochthonous populations along the distribution range of the species from 14 countries (six in Italy; three in Croatia and France; two in Germany, Poland, and Rumania; and one in Austria, Bulgaria, Czech Republic, Serbia, Slovak Republic, Slovenia, Spain, and Switzerland; Figure 1A and Table 1). In each population, leaves of 5(4) individuals were collected and preserved in silica gel (126 individuals in total). Genomic DNA was extracted from leaf material stored in silica gel using a DNA extraction kit (Qiagen DNeasy Plant Mini Kit) following the manufacturer’s instructions. The RAD library was prepared as in Balao et. al. (2020). Genomic DNA was digested by using high-fidelity *Sbf*I (New England Biolabs) and the resulting fragments were double-barcoded. The library was sequenced in a separate lane of an Illumina flowcell HiSeq 2500 at the VBCF NGS Unit (www.vbcf.ac.at/ngs) as 100 bp single-end reads.

### 2.2 RAD-seq catalogue building and SNP calling

Data processing was performed using the following workflow: (i) *Quality filtering and demultiplexing* of the library were performed with deML (Renaud et al., 2015) and STACKS ver. 2.55 (Rochette et al., 2019). Low-quality reads and reads without the restriction cut site were discarded. (ii) *SNP calling and genotyping*. Reads were mapped using Bowtie2 (Langmead & Salzberg, 2012) to the published *A. alba* draft genome (Mosca et al., 2019) using ––very-sensitive mode. Reads with mapping quality < 30 and coverage > 200 were discarded and we marked PCR duplicated with *samtools* 1.16.1 (Danecek et al., 2021). The genome-aligned reads were assembled into RAD loci using *ref_map.pl* code implemented through STACKS and the *populations* STACKS pipeline was used to export read data into various formats for subsequent analyses. We retained loci that exhibited a presence in a minimum of 70% of individuals throughout the entire dataset. In the subsequent analyses, we used 22019 SNPs dataset.

### 2.3 Population genetic diversity

Basic population genetic statistics were assessed and described using standard population genetic parameters. We estimated for each population in STACKS v2.55, the number of private alleles, number of polymorphic sites, expected heterozygosity (*H*_e_) and observed heterozygosity (*H*_o_), inbreeding coefficients (*F*_is_), and nucleotide diversity (π) with their standard errors (SE). In accordance with the geographical distribution of the populations under investigation, a systematic grouping approach was employed based on geographical regions of previous studies (Konnert & Bergmann, 1995; Piotti et al., 2017; Teodosiu et al., 2019). Populations located in the southeastern regions were designated as “S-C Apennines,” while those situated in the southwestern areas were grouped as “S Carpathians-Balkans.” Among the northern populations, the northwest was bifurcated into two distinct groups: “N Carpathians” and “E Alps. Meanwhile, the northeastern populations were similarly divided into “W Alps” and “N Balkans” (Figure 1A).

### 2.4 Phylogenomic analyses and genetic structure

A maximum-likelihood phylogeny was inferred using RAxML-ng v. 1.10 (Stamatakis, 2014). The analysis was conducted under the GTR+GAMMA model and Felsenstein’s model was used to address ascertainment bias in SNP data. To mitigate this bias, the +ASC_FELS parameter was employed, with a specific value (802204) denoting the invariable sites that were incorporated into the analysis. Branch support values were computed from 1000 bootstrap replicates. To situate the evolutionary position and as a point of reference for understanding the evolutionary relationships we introduced five individuals of *A. cephalonica* from different populations as outgroups of the phylogenetic tree (Balao et al., 2020).

To assess the genetic structure of different populations a principal component analysis (PCA) was estimated using the R package *adegenet* (Jombart & Ahmed, 2011). Additionally, we performed a Bayesian assignment analysis with ADMIXTURE version 1.3.0 (Alexander et al., 2015) using the 22019 SNP dataset. We conducted ADMIXTURE analysis by running the software with different levels of grouping (*K*), ranging from 1 to 10. For each level of grouping, we performed 100 replicates. Best-fitting *K* was chosen as the lower CV estimated.

### 2.5 Isolation-by-Distance (IBD) and geographical patterns of genetic diversity

The extent of genetic differentiation among various locations was quantified using the StAMPP (Pembleton et al., 2013). A matrix of *F*_ST_ scores was generated for each pair of populations under investigation. To examine the relationship between genetic disparities (represented by *F*_ST_) and geographical separation, we conducted an isolation by distance (IBD) analysis to explore the impact of geographic distance on the genetic differentiation across all 26 populations. This was accomplished by gauging the correlation between genetic differentiation and geographic distance through the utilization of the R package darT (Gruber et al., 2018). To investigate the potential relationship between genetic diversity estimators and geographical variables, we also formulated various geographically explicit generalized linear models (GLMs) using both genetic and geographical data. We considered the nucleotide diversity (π) and the number of private alleles as the dependent variables, while latitude and longitude as independent variables. The relationship between private alleles and geographic coordinates was analyzed using a Generalized Linear Model (GLM) with a Poisson distribution as the link function. In contrast, to assess the correlation between π and geographical coordinates, we applied a GLM with a Gaussian distribution.

### 2.6 Population splits and migration topology

TreeMix version 1.13 (Pickrell & Pritchard, 2012) was employed to analyze LD-pruned data, with the aim of inferring gene flow events among the 26 populations. For these analyses, we also removed missing data from the dataset, resulting in a 4821 SNPs dataset. In these analyses, we compared the likelihood from 1 to 5 migration events (m). Each analysis consisted of 500 bootstrap replicates with the population 20 used as outgroup. In addition, our analysis was extended to a regional scale by merging populations into four distinct regions: Southeast (SE), Southwest (SW), Northeast (NE), and Northwest (NW). The analysis was performed ten times across a range of migration event values (m) from 0 to 4, with each iteration supported by 500 bootstrap replicates. The SE population was consistently used as the outgroup across these analyses.

### 2.7 Demographic reconstructions

We studied the *A. alba* past demography, for the four lineages (SW, SE, NW and NE) obtained in the PCA and phylogenetic studies, by inferring the effective population size (*N*_e_) over time using the composite likelihood approach with a multi-epoch model implemented in the *Stairway plot* v2.1.1 software. We identified loci out of Hardy– Weinberg equilibrium using VCFTOOLS and discarded sites with evidence of heterozygote excess. We polarized the SNPs using a *A. cilicica* sample as outgroup. Then, we estimated the Site Frequency Spectrum (SFS) from 18512 SNPs by employing easySFS (accessible at https://github.com/isaacovercast/easySFS) to mitigate the impact of incomplete data and optimize the overall count of SNPs using a hypergeometric projection of the dataset. *Stairway plot* was run considering 25-years generation time and assuming a mutation rate of 1.23×10^−8^ per site per generation per locus (De La Torre et al., 2017; Sánchez-Robles et al., 2022). We used the recommended (nseq□−□2) ÷ 4 break points (where nseq is the number of haplotypes sampled) and 200 replicate runs for each population.

## 3 RESULTS

The 100 bp Illumina RAD-seq for 126 DNA samples (1.31 million reads on average per sample) representing all the 26 silver fir populations produced in the final STACKS catalogue a total of 779282 RAD loci with an average coverage per sample of 34.3 ± 14.2 (s.d.) reads per locus. We found a 6.62 % of missing RAD loci among samples with a mean of 1458.19 ± 63.81. After retaining only polymorphic RAD loci that were present in at least 88 individuals and had a maximum coverage of 100 reads per locus, we obtained a final data set with 8711 RAD loci and a total of 22019 SNPs.

### 3.1 Gene diversity

The analysis of genetic diversity (Table 2) showed that the southern populations exhibited the higher number of private alleles (518.57 ± 73.17). The highest average values of private alleles were found in the south-centre Apennines (661.33 ± 87.82) and the south Carpathians-Balkans (386.5 ± 87.65). S-C Apennines and S Carpathians-Balkans populations showed the highest values of nucleotide diversity, which ranged from 0.095 to 0.102. Specifically, populations of National Park Aspremonte and Laurenzana in the Italian Apennines (pops. 8-9), and Rila Monastery Nature Park in Bulgaria (pop. 21) showed a significant number of private alleles (828, 626, and 638, respectively; Table 2), indicating that these are probably more isolated than the other sampled locations.

**TABLE 2.** Genetic diversity of the 26 *Abies alba* populations analysed.

| Pop. | Polymorphic<br>alleles | Private<br>alleles | $H_o$ | $H_e$ | $\pi$ | $F_{IS}$ |
| --- | --- | --- | --- | --- | --- | --- |
| 1 | 62.9 | 332 | $0.087 \pm 0.001$ | $0.074 \pm 0.001$ | $0.085 \pm 0.001$ | $-0.002 \pm 0.005$ |
| 2 | 60.7 | 130 | $0.084 \pm 0.001$ | $0.073 \pm 0.001$ | $0.083 \pm 0.001$ | $-0.001 \pm 0.005$ |
| 3 | 68.2 | 223 | $0.097 \pm 0.001$ | $0.082 \pm 0.001$ | $0.094 \pm 0.001$ | $-0.007 \pm 0.005$ |
| 4 | 57.1 | 117 | $0.077 \pm 0.001$ | $0.070 \pm 0.001$ | $0.079 \pm 0.001$ | $0.004 \pm 0.004$ |
| 5 | 63.9 | 102 | $0.085 \pm 0.001$ | $0.076 \pm 0.001$ | $0.085 \pm 0.001$ | $0.002 \pm 0.003$ |
| 6 | 70.7 | 275 | $0.094 \pm 0.001$ | $0.082 \pm 0.001$ | $0.093 \pm 0.001$ | $-0.001 \pm 0.005$ |
| 7 | 63.0 | 530 | $0.095 \pm 0.001$ | $0.08 \pm 0.001$ | $0.095 \pm 0.001$ | $0.001 \pm 0.005$ |
| 8 | 68.8 | 626 | $0.099 \pm 0.001$ | $0.085 \pm 0.001$ | $0.101 \pm 0.001$ | $0.003 \pm 0.004$ |
| 9 | 77.2 | 828 | $0.094 \pm 0.001$ | $0.087 \pm 0.001$ | $0.100 \pm 0.001$ | $0.012 \pm 0.005$ |
| 10 | 61.8 | 115 | $0.087 \pm 0.001$ | $0.074 \pm 0.001$ | $0.084 \pm 0.001$ | $-0.006 \pm 0.004$ |
| 11 | 63.4 | 168 | $0.089 \pm 0.001$ | $0.076 \pm 0.001$ | $0.087 \pm 0.001$ | $-0.005 \pm 0.005$ |
| 12 | 59.5 | 89 | $0.077 \pm 0.001$ | $0.072 \pm 0.001$ | $0.081 \pm 0.001$ | $0.009 \pm 0.003$ |
| 13 | 62.6 | 147 | $0.084 \pm 0.001$ | $0.075 \pm 0.001$ | $0.085 \pm 0.001$ | $0.002 \pm 0.003$ |
| 14 | 61.0 | 167 | $0.077 \pm 0.001$ | $0.074 \pm 0.001$ | $0.083 \pm 0.001$ | $0.013 \pm 0.004$ |
| 15 | 67.1 | 418 | $0.091 \pm 0.001$ | $0.079 \pm 0.001$ | $0.089 \pm 0.001$ | $-0.001 \pm 0.004$ |
| 16 | 57.4 | 62 | $0.077 \pm 0.001$ | $0.071 \pm 0.001$ | $0.080 \pm 0.001$ | $0.008 \pm 0.003$ |
| 17 | 69.8 | 324 | $0.081 \pm 0.001$ | $0.081 \pm 0.001$ | $0.091 \pm 0.001$ | $0.023 \pm 0.004$ |
| 18 | 63.1 | 223 | $0.093 \pm 0.001$ | $0.078 \pm 0.001$ | $0.093 \pm 0.001$ | $0.000 \pm 0.004$ |
| 19 | 74.4 | 301 | $0.074 \pm 0.001$ | $0.084 \pm 0.001$ | $0.094 \pm 0.001$ | $0.045 \pm 0.004$ |
| 20 | 70.8 | 370 | $0.091 \pm 0.001$ | $0.083 \pm 0.001$ | $0.095 \pm 0.001$ | $0.008 \pm 0.006$ |
| 21 | 78.3 | 638 | $0.104 \pm 0.001$ | $0.089 \pm 0.001$ | $0.102 \pm 0.001$ | $-0.004 \pm 0.005$ |
| 22 | 65.8 | 346 | $0.099 \pm 0.001$ | $0.083 \pm 0.001$ | $0.098 \pm 0.001$ | $-0.001 \pm 0.004$ |
| 23 | 72.9 | 292 | $0.096 \pm 0.001$ | $0.084 \pm 0.001$ | $0.096 \pm 0.001$ | $0.000 \pm 0.005$ |
| 24 | 65.8 | 128 | $0.088 \pm 0.001$ | $0.076 \pm 0.001$ | $0.086 \pm 0.001$ | $-0.002 \pm 0.004$ |
| 25 | 66.0 | 138 | $0.089 \pm 0.001$ | $0.078 \pm 0.001$ | $0.088 \pm 0.001$ | $-0.001 \pm 0.004$ |
| 26 | 65.6 | 135 | $0.088 \pm 0.001$ | $0.077 \pm 0.001$ | $0.087 \pm 0.001$ | $-0.002 \pm 0.004$ |

In general, the northern populations showed a lower number of private alleles (189.16 ± 22.5). The individuals from the north Carpathians and east Alps showed a smaller average number of private alleles (162.43 ± 44.55) than the west Alps (178.78 ± 26.87; Table 2). Furthermore, the northern populations displayed a lower average genetic diversity (π = 0.87) in contrast to the southern populations (π = 0.098). The lower nucleotide diversity values (≤□0.081) were found in west and east Alps (Table S2). Individuals from Rauensteim in Germany (pop. 12) and Puščava in north Slovene Prealps (pop. 16) exhibited the least number of private alleles (62 and 89, respectively). The *F*_is_ values were homogeneous (from –0.007 to 0.045) and did not exhibit statistically significant differences when examined across population groups and geographic locations (Table 2).

### 3.2 Phylogenomic analyses and genetic structure

The PCA (Figure 1B) and phylogenomic analyses (Figure S1) demonstrated high congruence, revealing a clear geographic gradient both north/south and, to a lesser extent, east/west, which culminated in the delineation of four major genetic lineages (SW, SE, NW and NE). In PCA, the first principal component (PCA1), explaining the 59.7% of the variance, prominently distinguishes populations into two distinct regions: north and south. The second principal component (PCA2), which accounts for the 26.5% of the variance, delineates the separation between western and eastern populations, both in the northern and southern regions. In the phylogenetic tree, the first clade represents populations from southeastern (SE) range of the species and includes populations from S Balkans and the populations from S and SE Carpathians. The second grouping, which greatly differentiated from the other groups and resulted in the most related to the first one, represented the southwestern (SW) populations and includes the populations from S and C Apennine. The third major group clustered mainly the northeastern (NE) populations, and it was further subdivided into two well-differentiated subclades. The first subclade corresponded to populations from N Carpathians (pops. 24-26), and the second subclade includes the populations from E Alps (pops. 12 and 14-16). The fourth major group included populations from the W Alps and represents mostly all the individuals from continental populations of the northwestern (NW) range of the silver fir (pops. 1-6, 10-11 and 13). The phylogenetic tree (Figure S1) also showed and additional clade including the N Balkans populations and clustered the intermediate populations that we observed in the PCA (pops. 17-19).

### 3.3 Geographical patterns of diversity and Isolation-by-Distance (IBD)

Notably, the highest population pairwise-*F*_ST_ values were observed between the southern populations (from S-C Apennines and S Carpathians-Balkans) with NW (from W Alps) and NE (from E Alps and N Carpathians) populations (Figure 1C). In contrast, genetic differentiation among the southern populations appeared to be minimal. Interestingly, the three populations in the north Carpathians region exhibited nearly identical genetic profiles, suggesting a possible recent divergence event. Furthermore, we found a significant relationship between the genetic diversity estimators (π and private alleles) and latitude. The analysis clearly indicated that the southern regions, particularly the S-C Apennines and the S Carpathians-Balkans, demonstrated a significantly higher presence of private alleles (with *p*-values for private alleles and π both being < 0.001; see Figure 2A). Conversely, regions such as the N Carpathians and E Alps displayed a comparatively diminished presence of private alleles. Additionally, an inverse relationship was observed between latitude and the prevalence of private alleles; their frequency decreased northward across the geographical region. This trend was consistently mirrored in the genetic diversity values (Figure 2B), showing a decreasing gradient of genetic diversity with increasing latitude. In contrast, the southern regions consistently exhibited elevated genetic diversity values. Furthermore, GLM analyses also showed a significant relationship between genetic diversity estimators (π and private alleles) with longitude (for private alleles, Χ_2_ = 1866.1, *p*-value < 0.001 and for π, Χ_2_ = 16.662, *p*-value < 0.001; Figure S2), with a clear upward trend in both private alleles and genetic diversity (π) as populations move further eastward. Among these populations, those belonging to the S Carpathians-Balkans region exhibit the highest levels of genetic diversity and display a significant number of private alleles. Conversely, populations located farther west, such as those in the W Alps region, showed comparatively lower values. An exception to this trend was observed in the N Carpathians region, where, despite its eastern location, lower values were recorded.

**FIGURE 2.**
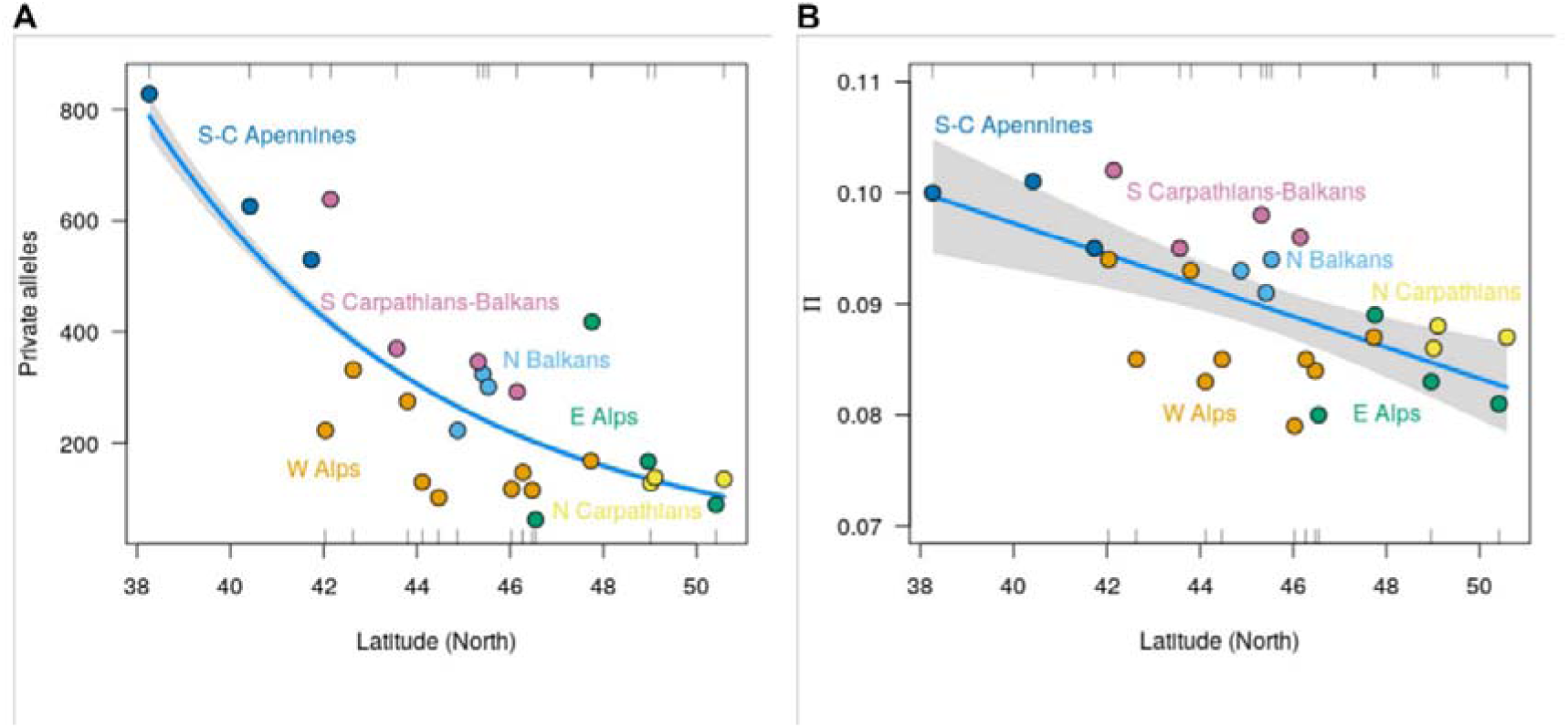
Latitudinal variation in genetic diversity of the *Abies alba* populations. A) private alleles and B) nucleotide diversity (π). Color denotes the main geographic areas.

### 3.4 Population coancestry and migration

Admixture results (Figure 3A), revealed that *K*=2 showed the best fit with the lowest CV. The best likelihood analysis, *K*=2, discerned again the southern and northern populations. The red cluster corresponded to all southern populations originating from the S-C Apennines and the S Carpathians-Balkans region. On the other hand, the blue cluster represented all populations located further north. In addition, we explored *K*=3 and *K*=4, which exhibited clear differentiation into four clusters, corresponding to both north-south and east-west divisions. The northern populations were further subdivided into the north-western populations from the Pyrenees, the western Alps in France, Germany and Switzerland (pops. 1, 2, 4 and 10-11), and the north-eastern populations from the eastern Alps in Germany, Czech Republic, Austria, and Slovenia, along with the eastern Carpathians populations from the Slovak Republic and Poland (pops. 12, 14-16 and 24-26). On the other hand, the southern populations were divided into two groups as well. The first group encompassed the southwest populations, including the previously mentioned S-C Apennines populations (pops. 7-9). The second group represented the southeast populations, corresponding to the S Carpathians-Balkans populations (pops. 20-23). However, some samples exhibited a high probability of genetic admixture, indicating the presence of mixed ancestry in multiple populations. The population from Corsica showed genetic influence from northwest populations, as well as that of the Dolomite Alps. While the Croatian populations of the Northern Balkans showed varying degrees of genetic influence. Individuals from population 19 exhibited a greater genetic influence from the south, however, individuals from populations 18 and 19 showed an influence from both, south and northwest lineages. The TreeMix results (Figure 3B) supported the same migration patterns between populations from different regions. The best model supported a unique migration event (m=1) from Romanian populations in S Carpathians to the N Carpathians populations.

**FIGURE 3.**
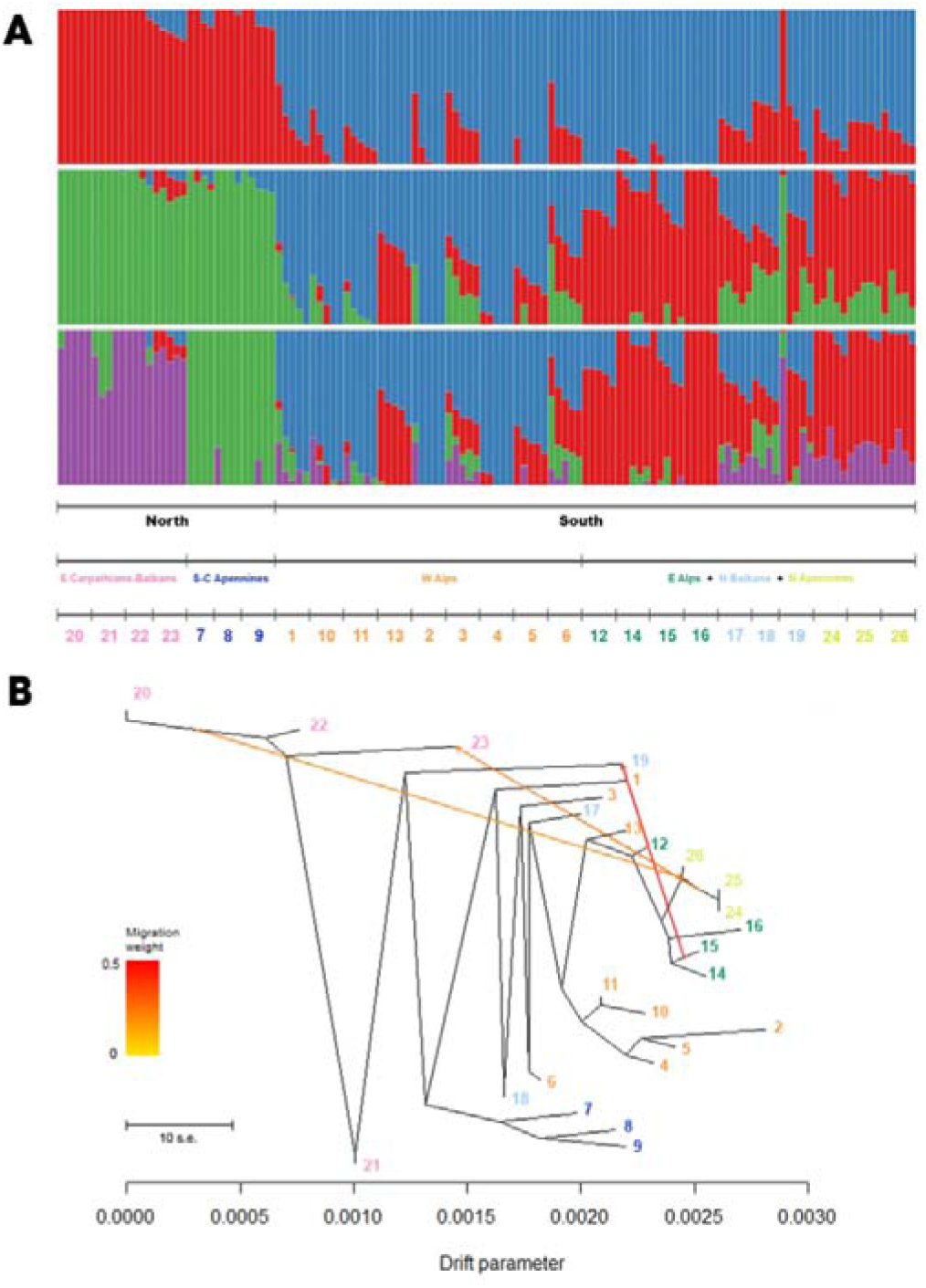
ADMIXTURE analysis reveals genetic clusters among 26 silver fir populations: *K*=1 separates them into northern and southern groups; *K*=3 splits the northern group into northeast and northwest; *K*=4 resolves four clusters—northeast, northwest, southeast, and southwest. B) Inferred principal migration events (m = 3) among the 26 populations using TreeMix. Migration weights are visualized by color intensity, with red indicating higher migration weights and yellow representing lower migration weights, elucidating the direction and relative magnitude of gene flow.

This introgression was also supported for m=2-5. In the model incorporating two migration events, it was additionally observed that the ancestral populations from Switzerland and Germany in the Western Alps (pops. 10-11) had interactions with other western Alps populations. Furthermore, m=3-5 analyses suggested introgression between Croatian populations in N Balkans and Serbian populations in S Carpathians-Balkans. A fourth putative migration (m=4-5) would occurred from Italian populations in S-C Apennines to Bulgarian populations in S Carpathians-Balkans. Lastly, we observed an additional introgression (m=5) between W Alps populations, specifically, Italian and Pyrenean populations.

### 3.5 Demographic reconstructions

The demographic analysis unveiled variations in the dynamics of effective population size (*N*_e_) over temporal scales between northern and southern regions (Figure 4). The northern regions were characterized by a significant population decline, with *N*_e_ decreasing notably from 1 Ma to 100,000 years ago, a trend that persisted until approximately 200,000 years ago. Around 50,000 years ago, these regions experienced a rapid exponential growth in *N*_e_, culminating in a state of population stability. Conversely, the southern populations did not undergo such a pronounced decline during the same period (Figure 4). They exhibited a gradual decline in population size over time, extending to the present day. Specifically, in the last 30,000 years, the southern regions have shown a continuous decline (Figure S3), with the rate of decrease notably intensifying approximately 8,000 years ago. In contrast, the demographic trajectories of the northern populations have been markedly different. These populations showed growth during the last glaciations, persisting until about 10,000 years ago, at which juncture a gradual decline began. Notably, this decline appears also to have accelerated from 8,000 years ago onwards.

**FIGURE 4.**
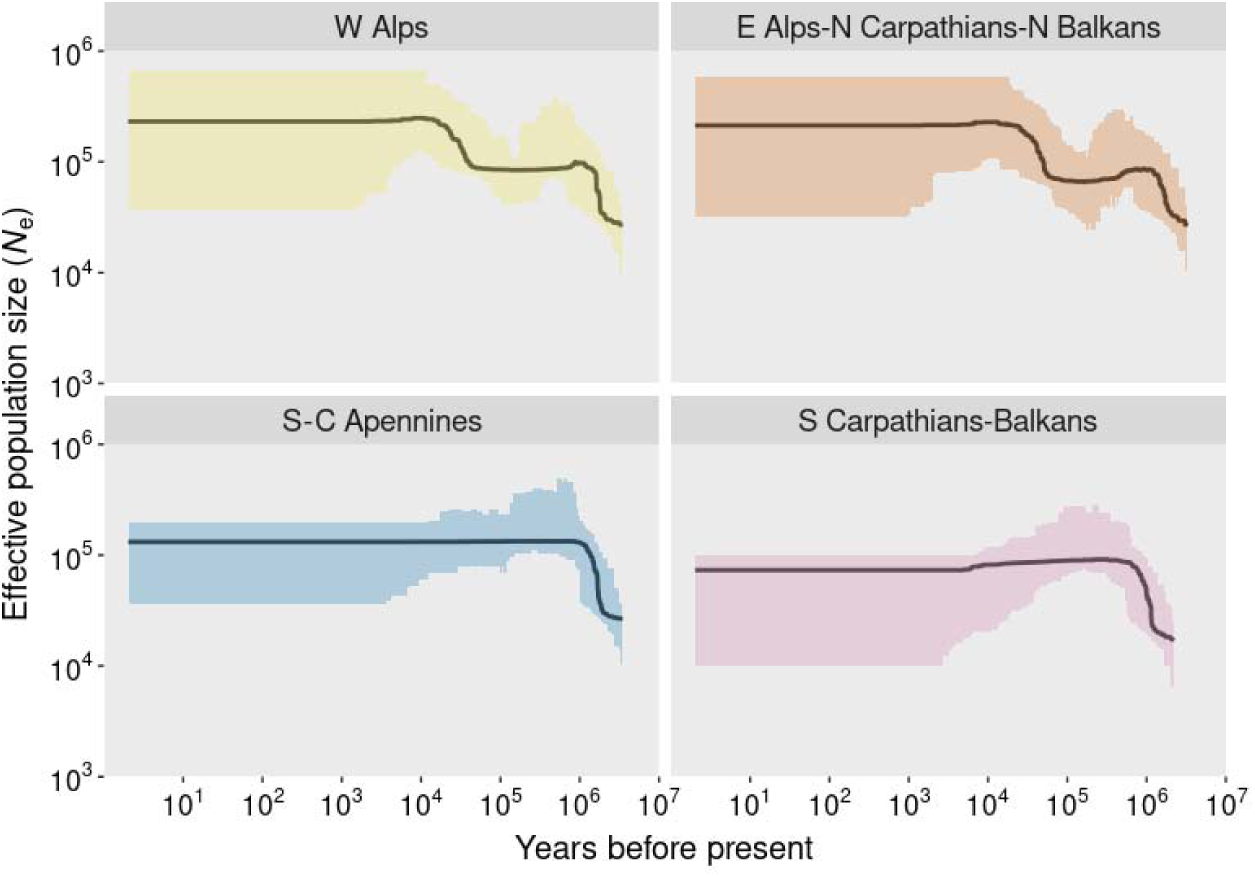
Demographic history of *Abies alba* populations via Stairway Plot illustrating changes in effective population size (*N*_e_) over the past 5 million years up to the present.

## 4 DISCUSSION

### 4.1 Landscape genetic structure

Based on our genome-wide analysis, four major phylogeographical lineages were detected, with a clear north/south differentiation and a moderately east/west geographic gradient. These findings not only are supported by multiple regional studies across the distribution of *A. alba* but also synthesize and unify the disparate pieces of evidence that have been emerging, providing a comprehensive biogeographic and genetic overview of the species. Regarding the Apennine populations, significant genetic divergence between the central and southern (the SW lineage) and those from the northern populations (NW lineage) was observed, as demonstrated by allozyme and microsatellite markers (Parducci et al., 1996; Piotti et al., 2017). This latter study not only pinpointed a genetic demarcation in the central Apennines, corresponding to pops. 5-9 which were part of their analysis, but also linked the divergent evolution of these groups over the last 400,000 years to both tectonic (Satolli & Calamita, 2008) and ecological factors (Combourieu-Nebout et al., 2015). Interestingly, our results highlight a historical trans-Adriatic connection between silver fir populations in the central and southern Apennines (SW lineage) and those in the Balkans and southern Carpathians (SE lineage). This connection is supported by genetic similarities across several molecular markers, as evidenced in previous studies (Liepelt et al., 2002; Longauer et al., 2003; Piotti et al., 2017). Also, Patacca et al. (2008) demonstrated that the Apulian platform (S Apennines) connected several times to the Balkans by a trans-Adriatic land bridge, additionally, the presence of shared haplotypes in the Balkan and Italian Peninsulas for other tree species such *Quercus cerris* has been found (Bagnoli et al., 2016). Notably, the SE lineage emerges as the most ancestral group in the phylogram, positioned closer to *A. cilicica* samples, which supports the hypothesis of an eastern origin for the species (Balao et al., 2020; Linares, 2011). Moreover, within the Carpathians, a distinct genetic differentiation is evident, segregating the S and SE populations (the SE lineage) from those in the NE and W Carpathians (NE lineage), a pattern also noted by Gömöry et al. (2012) using mitochondrial markers.

About the two genetic lineages of the northern range (NE and NW), all phylogenetic analyses corroborated the differentiation of the westernmost populations from Pyrenees, NW and SW Alps (continental French and Swiss populations), northwesternmost Apennine and that from S Black Forest, from those of the eastern part, clustering populations from NE and NW Carpathians, and also populations from SE and NE Alps, S Bohemia and Thuringian Forest. A parallel west-east geographical pattern was observed using mtDNA markers, pinpointing a contact zone between these two phylogeographical lineages in Croatia, Slovenia, and northeastern Italy, a finding that was similarly identified by Breitenbach-Dorfer et al. (1997). Additionally, the genetic analyses revealed a strong linkage between the western Emilian Apennine population (pop. 5) and those from the SW and NW Alps, as well as the S Black Forest, with all analyses clustering these groups together.

Regarding the origin of populations in the southwestern Alps, our findings align with those presented by Sagnard et al. (2002), who examined 16 populations from the French southwestern Alps (Sub-Mediterranean, Intermediate and Ligurian Alps). They observed no clear geographical structure among these populations, nor did they detect isolation-by-distance, implying significant historical gene flow between populations. This suggests that all the silver fir populations in the southwestern Alps likely originated from a common glacial refugium, which was probably situated in the Ligurian Apennines.

The placement of the Corsican population in our analysis revealed ambiguity. Phylogenetic analysis positioned it between the northern Balkans and Apennines, whereas PCA suggested closer affinity to the populations from the southwestern French Alps, southern Italian Alps, and the Pyrenees. This indeterminate positioning aligns with Fady et al. (1999), who observed that a population near our Corsican population 3 was relatively isolated but affiliated with the southwestern French Alps, suggesting shared glacial ancestry. Furthermore, palynological evidence from Reille et al. (1999) indicated that the migration of the species to the island occurred only during the late Holocene, adding another layer of complexity to its historical biogeography.

Finally, although certain studies had previously considered the Pyrenees population to be isolated since the last glacial period (Konnert & Bergmann, 1995), or even earlier (Scotti-Saintagne et al., 2021), it clearly clustered within the NW group (Figures 1B, 3A and S1). This relationship was corroborated by the genetic link between a Pyrenean population and a NW French Alps population, identical to pop. 4, as described in Fady et al. (1999).

### 4.2 Legacies of the Holocene colonization

The genetic diversity patterns (private alleles and π) and the phylogenomic and genetic structure analyses (Figures 1B and 3A) suggested that silver fir survived in different southern refugia area during the last glaciations. Southern populations had higher diversity and private alleles that northern ones. Our data strongly confirm the existence of a glacial refugium in the southern Apennines (most probably in Aspromonte, southern Calabrian Apennines), which appears to have remained isolated from the northern Apennines refugia. Consequently, these populations experienced limited expansions and have remained genetically distinct from those in the northern Apennine chain (see also the following section on demography). Congruently, numerous studies (Bergmann et al., 1990; Konnert & Bergmann, 1995; Liepelt et al., 2009) have supported the existence of this refugium in southern Italy, confirming the disconnection of the Calabrian silver fir from those of northern regions since the Last Glacial Maximum. Whereas the existence of effective glacial refugium of silver fir in the southern or south-eastern Balkan is undoubtedly given our results (populations with high gene diversity and private alleles), its position has not been precisely assigned previously due to a lack of fossil records (Liepelt et al., 2009), although, our genetic results clearly point to Rila-Rhodope Massif (pop. 21). However, due to the genetic relationship of the W and NE Carpathians populations, another refugium from north Balkan Peninsula (most likely in N Croatia) cannot be discarded. Lastly, based on the intermediate position of the north Balkans populations (pops. 17-19; Figures 1B, 3A and S1), we propose that silver fir from Apennine and Balkan refugia had met and developed in two introgression zones, one in northwestern and northeastern Croatia and probably in the foothills of the Slovenian Alps, and a second in High Tatra, Slovak Carpathians, where silver fir from the southern Balkan area had met silver fir from northern Apennine during its migration.

Our data strongly confirmed the source of the NW genetic lineage, from SW genetic lineages whereas; the NE genetic lineage arrive from East Alpine populations. These results robustly corroborate the hypotheses regarding the primary routes and subroutes for the postglacial recolonization of northwestern, central, and northeastern European regions by the silver fir. We found strong evidence supporting both the north Apennine and southern Balkan Peninsula as key origins of recolonization. Moreover, the subdivision of these main routes into the “West Alpine” and “East Alpine” routes from the Apennines, as well as the “West Balkan” and “East Balkan” routes from the Balkan Peninsula, has also been validated by our findings. This confirmation extends to the intricate pathways described, from the Ligurian mountains that outflanking the Alps. Similarly, our data substantiate the movement of populations along the Dinaric Alps towards the eastern Alps and through Southern Bulgaria into the Carpathians. These results not only validate the proposed historical migration patterns (see Karl, 1980; Konnert & Bergmann, 1995; Liepelt et al., 2002, 2009) but also enrich our phylogenetic and biogeographic understanding of *A. alba* across its European range.

This complex biogeographic history contrast to some similar tree species, such spruce (*Picea* sp.) in North America, whose range shifted entirely since the last ice age (Williams et al., 2004). Here, relatively stable refugia suggest that some silver fir populations have persisted at suitable sites across Quaternary climatic oscillations until present, providing an outstanding biogeographic and conservation value (Hampe & Petit, 2005; Tzedakis et al., 2002). Furthermore, some of these relict silver fir populations, such those from southern Apennines, might have not been the source of major postglacial recolonizations, thereby preserving high genetic distinctiveness.

### 4.3 Conservation setting from demography and vulnerability to ongoing climate change

Although the demographic events that shaped the current spatial structure of genetic diversity of silver fir go back to the Pleistocene, human pressure during the Holocene likely led to an abrupt range decline, while recent drought events have caused widespread decreasing productivity and dieback of silver fir forest, mainly over its southernmost range limits.

Pleistocene glacial periods likely led to significant fluctuations in the demography of *A. alba* populations, contingent upon their geographical location within Europe (Scotti et al., 2023). In our study, populations from southern regions showed a comparatively stable trajectory over the years, acting as glacial refuges (Linares et al., 2011). These regions have exhibited a consistent ascendant trend from the early Pleistocene over a million years ago to the late Pleistocene approximately 200,000 years ago. Other authors also situated the demographic events that shaped the current spatial structure of genetic diversity of silver fir during the Middle Pleistocene (Scotti-Saintagne et al., 2021). Populations in the southern regions began stabilizing their growth to the present day, with an observable continuous decline during the last 30,000 years. Significant anthropogenic pressures would have impacted *A. alba* forests causing the rapid demographic decline from the last 8,000 years (Linares et al., 2011; Pons & Quézel, 1985) as occurred in *A. pinsapo* in south Iberian Peninsula (Sánchez-Robles et al., 2022). The dynamics of those populations that inhabit the southernmost margins of the silver fir distribution range might be critically important in determining the adaptive capacity of this tree to expected climate change (Hampe & Petit, 2005). Thus, selection for local adaptation might be expected in these marginal populations, as well as currently reduced gene flow, resulting in potential distinct ecotypes (Alberto et al., 2013). Notably, a distinct pattern is evident in northern regions, where populations that had been expanding for over a million year began experiencing a reduction in population size during the middle Pleistocene (from approximately a million years ago to around 200,000 years ago) due to glaciations in this era (Linares et al., 2011; Liepelt et al., 2009).

Northern populations would undergo exponential growth after the last glacial period, aligning with the colonization and expansion from glacial refugia of southern populations migrating northward at the late Pleistocene and the onset of the Holocene (Liepelt et al., 2002, 2009; Litkowiec et al., 2016; Terhürne-Berson et al., 2004). As the southern regions, the northern populations also suffered an abrupt decline about 8,000 years ago, which would correspond with the impact of human pressures. Hence, species distribution models suggest that the current distribution range of silver fir was determined by historical land use, confusing to some extent the potential impacts of climate change (Di Pasquale et al., 2014; Svenning & Skov, 2004; Tinner et al., 2013).

The current patterns of genetic diversity and populations dynamics provide valuable information to evaluate the risk of losing silver fir populations or habitats, that is, the vulnerability of different locations within the whole species distribution range (see also Tinner et al., 2013). Our demography inferences support that southern *A. alba* populations acted as glacial refuges for over a million years but recently suffered a declined due to human impact (Di Pasquale et al., 2014), while northern populations fluctuated with glaciations and colonization events (Svenning et al., 2008). As we noted above, the presence of local adaptations in those populations, that inhabit the rear edge of the silver fir distribution range, might be critical in determining the fate of this tree under a climate change scenario (Castagneri et al., 2014; Sánchez-Salguero et al., 2017). Tree species dominant in temperate forests of Europe, such as silver fir, reach their southern limit in the Mediterranean region, a climate change hotspot, where warming is enhancing aridity in the last decades at unprecedented rates (Luterbacher et al., 2012).

Indeed, recent drought events have caused widespread dieback and mortality in some of the main silver fir populations identified here as glacial refuges. Hence, dendroecological studies evidenced contrasting growth variability and climate sensitivity in *A. alba* over the distribution range, that are in line with post-glacial phylogeny and genetic diversity (Bosela et al., 2016).. Specifically, the growth of silver fir increased considerably during the last decades across central Europe (Buntgen et al., 2014), whereas there was a significant growth decline toward the Mediterranean ecotone in southern France and the Pyrenees (Cailleret et al., 2014; Linares & Camarero, 2012; Macias et al., 2006) and south-eastern European mountains in Slovenia, Croatia, Slovakia and Bosnia-Herzegovina (Diaci et al., 2011), according to glacial refugia identified here for the Balkans. Furthermore, different growth patterns have been also reported comparing northern and southern populations of silver fir in Italy (Carrer et al., 2010), according to the glacial refugium identified here for the Apennines.

Borrowing from the several studies reporting current and expected silver fir forests dieback in the main regions identified here as acting as glacial refuges, we conclude that those populations deserve priority conservation efforts, as they are disproportionately important for the long-term conservation of genetic diversity, phylogenetic history and evolutionary potential of silver fir. Hence, we demonstrate that the populations from the rear edges of the *A. alba* distribution range harbor higher diversity and private alleles, while they are currently in climate change hotspots, where drier and warmer summer conditions will represent the major climatic constraints to the persistence of silver fir.

## ACKNOWLEDGEMENTS

We would like to thank the Herbarium and Biology Services (CITIUS2) and the Informatic Service (SIC) from the University of Seville for the access to the facilities and bioinformatics resources.

## DATA AVAILABILITY STATEMENT

All data generated during this study are included in this published article and its supplementary information files. The raw Illumina sequencing data will be deposited on NCBI SRA (GenBank BioProject PRJNAXXXX, accessions YYYY) before the paper will be accepted for publication. Datasets and the R code used for analysis are available from GitHub (https://github.com/fbalao/albams).

## Supplementary information

**Figure S1.**
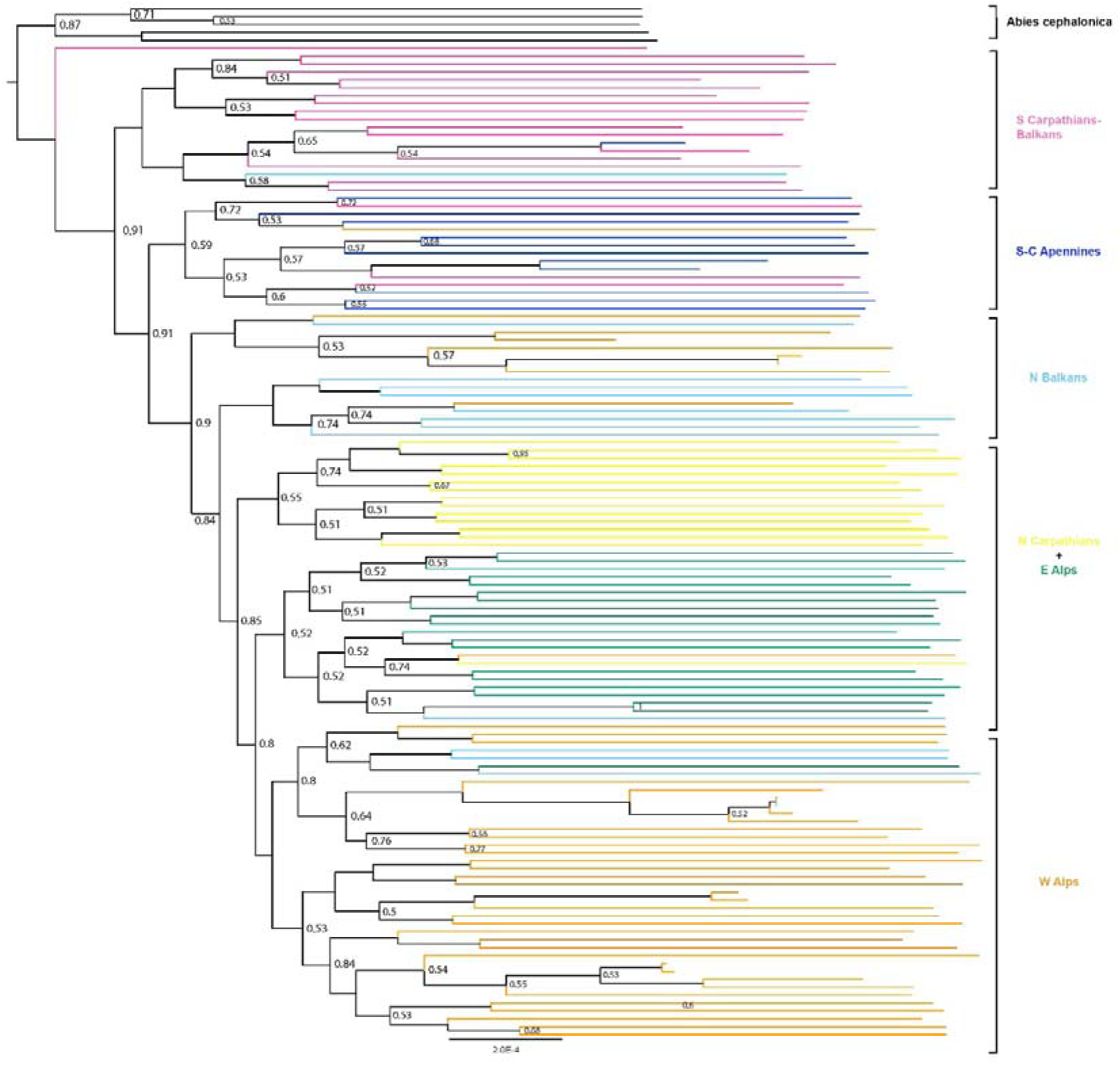
Phylogenetic tree of 126 tree samples from 26 Abies alba populations, constructed via maximum likelihood using RAxML-ng. Colors denote main geographical regions.

**Figure S2.**
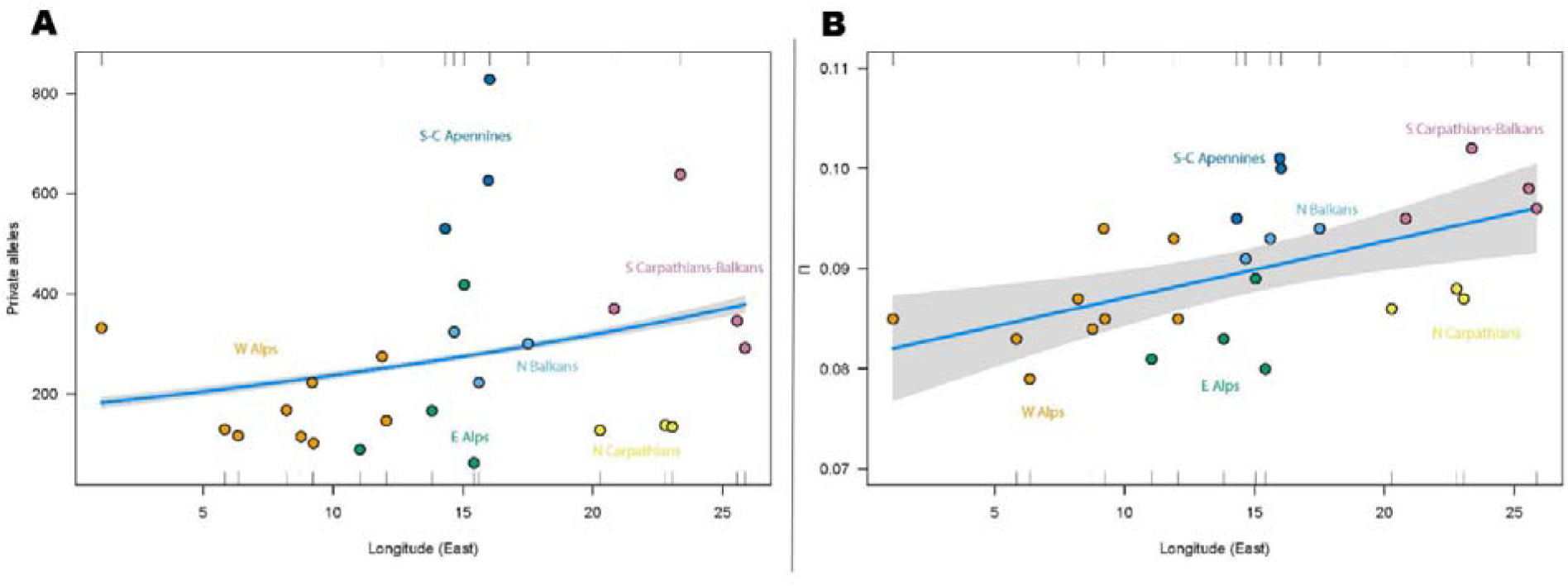
GLM analysis of genetic diversity vs. longitude in *A. alba* populations: (A) private alleles and (B) nucleotide diversity (π).

**Figure S3.**
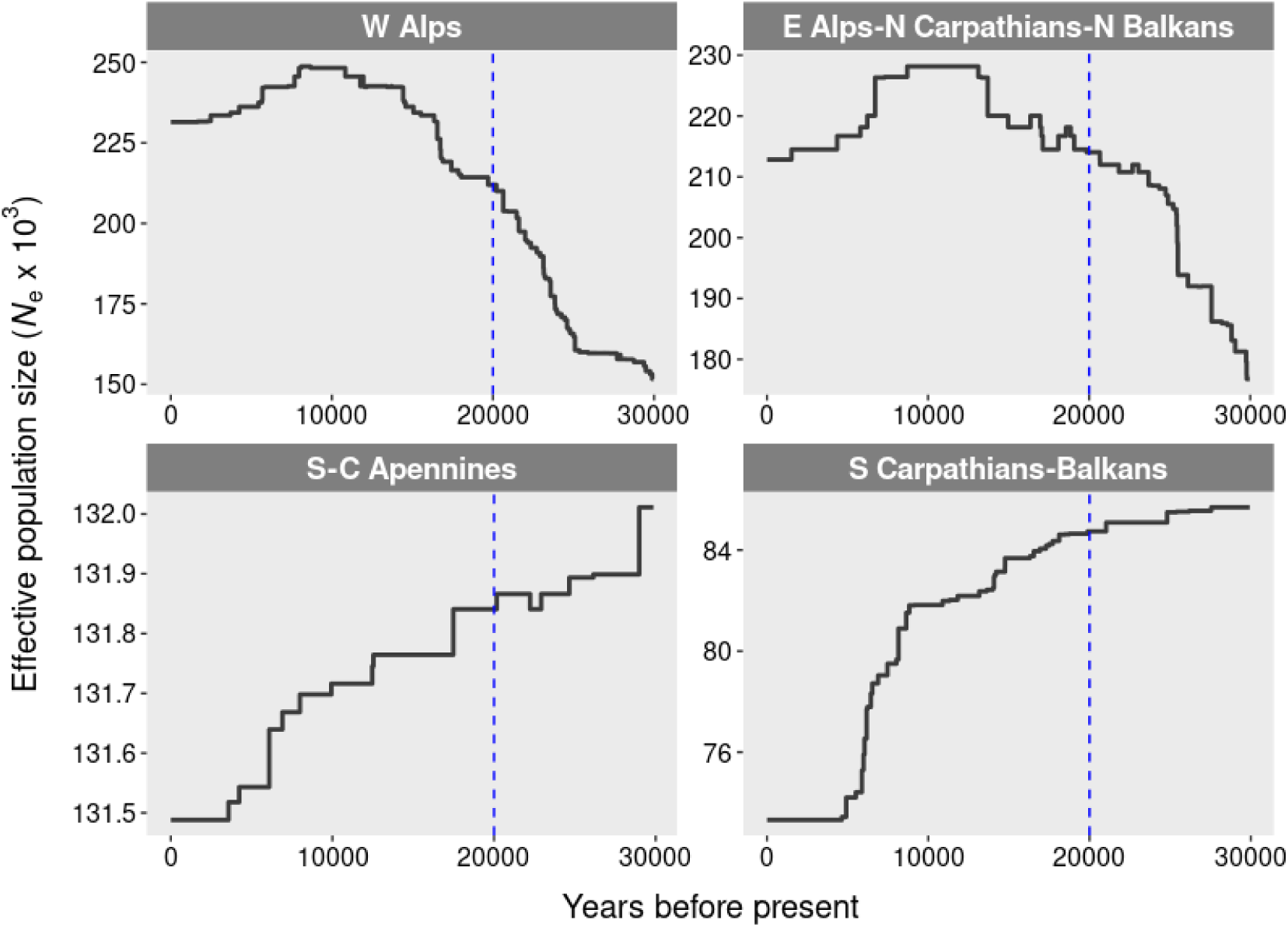
Stairway plot of *Abies alba* demographic history showing changes in effective population size (*N_e_*) over the last 30,000 years.

## REFERENCES

1. Aitken, S. N., Yeaman, S., Holliday, J. A., Wang, T. L., & Curtis-McLane, S. (2008). Adaptation, migration or extirpation: climate change outcomes for tree populations. Evolutionary Applications, 1, 95–111. 10.1111/j.1752-4571.2007.00013.x

2. Alberto, F.J., Aitken, S.N., Alía, R., González-Martínez, S.C., Hänninen, H., Kremer, A., Lefèvre, F., Lenormand, T., Yeaman, S., Whetten, R. and Savolainen, O. (2013). Potential for evolutionary responses to climate change – evidence from tree populations. Global Change Biology, 19, 1645–1661. 10.1111/gcb.12181

3. Alexander, D.H., Shringarpure, S.S., Novembre, J., & Lange, K.L. (2015). Admixture 1.3 Software Manual.

4. Allen, C. D., Breshears, D. D., & McDowell, N. G. (2015). On Underestimation of Global Vulnerability to Tree Mortality and Forest Die-off from Hotter Drought in the Anthropocene. Ecosphere, 6(8), 1–55. 10.1890/ES15-00203.1

5. Bagnoli, F., Tsuda, Y., Fineschi, S., Bruschi, P., Magri, D., Zhelev, P., Paule, L., Mc, S., González-Martínez, S., & Vendramin, G. G. (2016). Combining molecular and fossil data to infer demographic history of *Quercus cerris*: Insights on European Eastern Glacial refugia. Journal of Biogeography, 43(4), 679–690. 10.1111/jbi.12673

6. Balao, F., Lorenzo, M. T. C., Sánchez-Robles, J. M., Paun, O., García-Castaño, J. L., & Terrab, A. (2020). Early diversification and permeable species boundaries in the Mediterranean Firs. Annals of Botany, 125(3), 495–507. 10.1093/aob/mcz186

7. Barrett, R., & Schluter, D. (2008). Adaptation from Standing Genetic Variation. Trends in Ecology & Evolution, 23 (1), 38–44. 10.1016/j.tree.2007.09.008

8. Bergmann, F., Gregorius, H., & Larsen, J. F. (1990). Levels of genetic variation in European silver FIR (*Abies alba*). Genetica, 82(1), 1–10. 10.1007/BF00057667

9. Bosela, M., Popa, I., Gömöry, D., Longauer, R., Tobin, B., Kyncl, J., Kyncl, T., Nechita, C., Petráš, R., Sidor, C.G., Šebeň, V. & Büntgen, U. (2016), Effects of post-glacial phylogeny and genetic diversity on the growth variability and climate sensitivity of European silver fir. Journal of Ecology, 104, 716–724. 10.1111/1365-2745.12561

10. Breitenbach-Dorfer, M., Konnert, M., Pinsker, W., Starlinger, F., & Geburek, T. (1997). The contact zone between two migration routes of silver fir, *Abies alba* (Pinaceae), revealed by allozyme studies. Plant Systematics and Evolution, 206(1-4), 259–272. 10.1007/BF00987951

11. Büntgen, U., Tegel, W., Kaplan, J.O., Schaub, M., Hagedorn, F., Bürgi, M., Brázdil, R., Helle, G., Carrer, M., Heussner, K.-U., Hofmann, J., Kontic, R., Kyncl, T., Kyncl, J., Camarero, J.J., Tinner, W., Esper, J. & Liebhold, A. (2014). Placing unprecedented recent fir growth in a European-wide and Holocene-long context. Frontiers in Ecology and the Environment, 12, 100–106. 10.1890/130089

12. Cailleret, M., Nourtier, M., Amm, A., Durand-Gillmann, M., & Davi, H. (2014). Drought-induced decline and mortality of silver fir differ among three sites in Southern France. Annals of Forest Science, 71, 643–657. 10.1007/s13595-013-0265-0

13. Carrer, M., Nola, P., Motta, R. & Urbinati, R. (2010). Contrasting tree-ring growth to climate responses of *Abies alba* toward the southern limit of its distribution area. Oikos, 119, 1515–1525. 10.1111/j.1600-0706.2010.18293.x

14. Castagneri, D., Nola, P., Motta, R., & Carrer, M. (2014). Summer climate variability over the last 250 years differently affected tree species radial growth in a mesic Fagus–Abies–Picea old-growth forest. Forest Ecology and Management, 320, 21–29.

15. Cheddadi, R., Birks, H. J. B., Tarroso, P., Liepelt, S., Gömöry, D., Dullinger, S., Meier, E. S., Hülber, K., Maiorano, L., & Laborde, H. (2014). Revisiting tree-migration rates: *Abies alba* (Mill.), a case study. Vegetation History and Archaeobotany, 23(2), 113–122.

16. Combourieu-Nebout, N., Bertini, A., Russo-Ermolli, E., Peyron, O., Klotz, S., Montade, V., … & Sadori, L. (2015). Climate changes in the central Mediterranean and Italian vegetation dynamics since the Pliocene. Review of Palaeobotany and Palynology, 218, 127–147.

17. Danecek, P., Bonfield, J. K., Liddle, J., Marshall, J., Ohan, V., Pollard, M. O., Whitwham, A., Keane, T., McCarthy, S. A., Davies, R. M., & Li, H. (2021). Twelve years of SAMtools and BCFtools. GigaScience, 10(2), giab008.

18. Davis, M. B., & Shaw, R. G. (2001). Range shifts and adaptive responses to quaternary climate changes. Science, 292, 673–679.

19. De La Torre, A. R., Li, Z., Van De Peer, Y., & Ingvarsson, P. K. (2017). Contrasting Rates of Molecular Evolution and Patterns of Selection among Gymnosperms and Flowering Plants. Molecular Biology and Evolution, 34(6), 1363–1377. 10.1093/molbev/msx069

20. Diaci, J., Rozenbergar, D., Anic, I., Mikac, S., Saniga, M., Kucbel, S., Visnjic, C. & Ballian, D. (2011). Structural dynamics and synchronous silver fir decline in mixed old-growth mountain forests in Eastern and Southeastern Europe. Forestry, 84, 479–491.

21. Di Pasquale, G., Allevato, E., Cocchiararo, A., Moser, D., Pacciarelli, M. & Saracino, A. (2014). Late Holocene persistence of *Abies alba* in low-mid altitude deciduous forests of central and southern Italy: new perspectives from charcoal data. Journal of Vegetation Science, 25, 1299–1310.

22. *EUFORGEN*. (2011). European Forest Genetic Resources Programme. (s. f.). https://www.euforgen.org/

23. Fady, B., Forest, I., Hochu, I., Ribiollet, A. D., De Beaulieu, J. L., & Pastuszka, P. (1999). Genetic differentiation in *Abies alba* Mill. populations from south-eastern France. Forest Genetics, 6, 129–138.

24. Fady, B., Esposito, E., Abulaila, K., Aleksic, J. M., Alia, R., Alizoti, P., … & Westergren, M. (2022). Forest genetics research in the Mediterranean Basin: bibliometric analysis, knowledge gaps, and perspectives. Current Forestry Reports, 8(3), 277–298.

25. Farjon, A., & Rushforth, K. (1989). A classification of *Abies* Miller (Pinaceae). Notes of the Royal Botanical Garden of Edinburgh, 46(1), 59–79.

26. Gömöry, D., Paule, L., Krajmerová, D., Romšáková, I., & Longauer, R. (2012). Admixture of genetic lineages of different glacial origin: a case study of *Abies alba* mill. in the Carpathians. Plant Systematics and Evolution, 298(4), 703–712.

27. Gruber, B., Unmack, P. J., Berry, O. F., & Georges, A. (2018). dartr: An r package to facilitate analysis of SNP data generated from reduced representation genome sequencing. Molecular Ecology Resources, 18(3), 691–699.

28. Hampe, A., & Petit, R. J. (2005), Conserving biodiversity under climate change: the rear edge matters. Ecology Letters, 8, 461–467. 10.1111/j.1461-0248.2005.00739.x.

29. Hewitt, G. M. (1999). Post-glacial re-colonization of European biota. Biological Journal of the Linnean Society, 68, 87–112. 10.1111/j.1095-8312.1999.tb01160.x

30. Hewitt, G.M. (2000). The genetic legacy of the ice ages. Nature, 405, 907–913.

31. Hewitt, G. M. (2004). Genetic consequences of climatic oscillations in the Quaternary. Philosophical Transactions of the Royal Society of London. Series B: Biological Sciences, 359(1442), 183–195.

32. Hussendörfer, E. (1999). Genetic variation of silver fir populations (*Abies alba* Mill.) in Switzerland. Forest Genetics, 6(2), 101–113.

33. Jombart, T., & Ahmed, I. (2011). *Adegenet 1.3-1*: New tools for the analysis of genome-wide SNP data. Bioinformatics, 27(21), 3070–3071.

34. Kempf, M., Zarek, M., & Paluch, J. (2020). The pattern of genetic variation, survival and growth in the *Abies alba* Mill. population within the introgression zone of two refugial lineages in the Carpathians. Forests, 11(8), 849.

35. Keppel, G., Van Niel, K. P., Wardell-Johnson, G. W., Yates, C. J., Byrne, M., Mucina, L., … & Franklin, S. E. (2012). Refugia: identifying and understanding safe havens for biodiversity under climate change. Global Ecology and Biogeography, 21(4), 393–404.

36. Konnert, M., & Bergmann, F. (1995). The geographical distribution of genetic variation of silver fir (*Abies alba*, Pinaceae) in relation to its migration history. Plant Systematics and Evolution, 196(1-2), 19–30.

37. Karl, F. (1980). Waldgeschichtliche Grundlagen für die Ausscheidung von Ökotypen bei *Abies alba*. – Proceedings 3. IUFRO Tannensymposium Wien, pp. 158–168. – Wien: Österr. Agrar-Verlag.

38. Langmead, B., & Salzberg, S. L. (2012). Fast gapped-read alignment with Bowtie 2. Nature Methods, 9(4), 357–359.

39. Linares, J. C. (2011). Biogeography and evolution of *Abies* (Pinaceae) in the Mediterranean Basin: the roles of long-term climatic change and glacial refugia. Journal of Biogeography, 38(4), 619–630.

40. Linares, J. C., & Camarero, J. J. (2012). Growth patterns and sensitivity to climate predict silver fir decline in the Spanish Pyrenees. European Journal of Forest Research, 131, 1001–1012.

41. Linares, J. C., Delgado-Huertas, A., & Carreira, J. A. (2011). Climatic trends and different drought adaptive capacity and vulnerability in a mixed *Abies pinsapo*– *Pinus halepensis* forest. Climatic change, 105(1), 67–90.

42. Liepelt, S., Bialozyt, R., & Ziegenhagen, B. (2002). Wind-dispersed pollen mediates postglacial gene flow among refugia. Proceedings of the National Academy of Sciences of the United States of America, 99(22), 14590–14594.

43. Liepelt, S., Cheddadi, R., De Beaulieu, J., Fady, B., Gömöry, D., Hussendörfer, E., Konnert, M., Litt, T., Longauer, R., Terhürne-Berson, R., & Ziegenhagen, B. (2009). Postglacial range expansion and its genetic imprints in *Abies alba* (Mill.) — A synthesis from palaeobotanic and genetic data. Review of Palaeobotany and Palynology, 153(1–2), 139–149.

44. Litkowiec, M., Lewandowski, A., & Rączka, G. (2016). Spatial Pattern of the Mitochondrial and Chloroplast Genetic Variation in Poland as a Result of the Migration of *Abies alba* Mill. from Different Glacial Refugia. Forests, 7(12), 284.

45. Longauer, R. (2001). Genetic variation of European silver fir (*Abies alba* Mill.) in the Western Carpathians. Journal of Forest Science, 47(10), 429–438.

46. Longauer, R., Paule, L., & Andonoski, A. (2003). Genetic diversity of southern populations of *Abies alba* Mill. Forest Genetics, 10, 1–10.

47. Luterbacher, J., García-Herrera, R., Allan, A. R., Alvarez-Castro, B. G., Benito, G., Booth, J., … & Zorita, E. (2012). A review of 2000 years of paleoclimatic evidence in the Mediterranean. The Climate of the Mediterranean Region From the past to the future, 87–185.

48. Macias, M., Andreu, L., Bosch, O., Camarero, J.J. & Gutiérrez, E. (2006). Increasing aridity is enhancing silver fir (*Abies alba* Mill.) water stress in its south-western distribution limit. Climatic Change, 79, 289–313.

49. McDowell, N. G., Allen, C. D., Anderson-Teixeira, K., Aukema, B. H., Bond-Lamberty, B., Chini, L., … & Xu, C. (2020). Pervasive shifts in forest dynamics in a changing world. Science, 368(6494), eaaz9463. 10.1126/science.aaz9463.

50. Médail, F., & Diadema, K. (2009). Glacial refugia influence plant diversity patterns in the Mediterranean Basin. Journal of Biogeography, 36(7), 1333–1345.

51. Mosca, E., Cruz, F., Gómez-Garrido, J., Bianco, L., Rellstab, C., Brodbeck, S., Csilléry, K., Fady, B., Fladung, M., Fussi, B., Gömöry, D., González-Martínez, S. C., Grivet, D., Gut, M., Hansen, J. K., Heer, K., Kaya, Z., Krutovsky, K. V., Kersten, B., Liepelt, S., Opgenoorth, L., Sperisen, C., Ullrich, K.K., Vendramin, G.G., Westergren, M., Ziegenhagen, B., Alioto, T., Gugerli, F., Heinze, B., Höhn, M., Troggio, & Neale, D. B. (2019). A Reference Genome Sequence for the European Silver Fir (Abies alba Mill.): A Community-Generated Genomic Resource. G3: Genes, Genomes, Genetics, 9(7), 2039–2049.

52. Parducci, L., Szmidt, A. E., Villani, F., Wang, X. R., & Cherubini, M. (1996). Genetic variation of *Abies alba* in Italy. Hereditas, 125(1), 11–18.

53. Patacca, E., Scandone, P., & Mazza, P. (2008). Oligocene migration path for Apulia macromammals: the Central-Adriatic bridge. Bollettino Della Societa Geologica Italiana, 127(3), 337–355.

54. Pembleton, L. W., Cogan, N. O. I., & Forster, J. W. (2013). StAMPP: an R package for calculation of genetic differentiation and structure of mixed-ploidy level populations. Molecular Ecology Resources, 13(5), 946–952.

55. Pickrell, J. K., & Pritchard, J. K. (2012). Inference of population splits and mixtures from genome-wide allele frequency data. PLoS genetics, 8(11), e1002967.

56. Piotti, A., Leonarduzzi, C., Postolache, D., Bagnoli, F., Spanu, I., Brousseau, L., Urbinati, C., Leonardi, S., & Vendramin, G. G. (2017). Unexpected scenarios from Mediterranean refugial areas: disentangling complex demographic dynamics along the Apennine distribution of silver fir. Journal of Biogeography, 44(7), 1547–1558.

57. Pons, A., & Quezel, P. (1985). history of the flora and vegetation and past and present human disturbance in the Mediterranean region. Geobotany.

58. Reille, M., & Lowe, J. J. (1993). A re-evaluation of the vegetation history of the eastern Pyrenees (France) from the end of the last glacial to the present. Quaternary Science Reviews, 12(1), 47–77.

59. Reille, M., Gamisans, J., Andrieu-Ponel, V., & De Beaulieu, J. (1999). The Holocene at Lac de Creno, Corsica, France: a key site for the whole island. New Phytologist, 141(2), 291–307.

60. Renaud, G., Stenzel, U., Maričić, T., Wiebe, V., & Kelso, J. (2015). deML: robust demultiplexing of Illumina sequences using a likelihood-based approach. Bioinformatics, 31(5), 770–772.

61. Rochette, N. C., Rivera-Colón, A. G., & Catchen, J. M. (2019). Stacks 2: Analytical methods for paired-end sequencing improve RADseq-based population genomics. Molecular Ecology, 28(21), 4737–4754.

62. Sagnard, F., Barberot, C., & Fady, B. (2002). Structure of genetic diversity in *Abies alba* Mill. from southwestern Alps: multivariate analysis of adaptive and non-adaptive traits for conservation in France. Forest Ecology and Management, 157(1-3), 175–189.

63. Sánchez-Robles, J. M., García-Castaño, J. L., Balao, F., García, C., Terrab, A., & Lozano, S. T. (2022). Anthropogenic deforestation and climate dryness as drivers of demographic decline and genetic erosion in the southernmost European fir forests. European Journal of Forest Research, 141(4), 649–663. 10.1007/s10342-022-01467-3

64. Sánchez-Salguero, R., Camarero, J. J., Carrer, M., Gutiérrez, E., Alla, A. Q., Andreu-Hayles, L., Hevia, A., Koutavas, A., Martínez-Sancho, E., Nola, P., Papadopoulos, A., Pasho, E., Toromani, E., Carreira, J. A., & Linares, J. C. (2017). Climate extremes and predicted warming threaten Mediterranean Holocene firs forests refugia. Proceedings of the National Academy of Sciences, 114(47), E10142–E10150.

65. Satolli, S., & Calamita, F. (2008). Differences and similarities between the central and the southern Apennines (Italy): Examining the Gran Sasso versus the Matese-Frosolone salients using paleomagnetic, geological, and structural data. Journal of Geophysical Research, 113(B10).

66. Scotti, I., Lalagüe, H., Oddou-Muratorio, S., Scotti-Saintagne, C., Ruiz Daniels, R., Grivet, D., … & Vendramin, G. G. (2023). Common microgeographical selection patterns revealed in four European conifers. Molecular Ecology, 32(2), 393–411.

67. Scotti-Saintagne, C., Boivin, T., Suez, M., Musch, B., Scotti, I., & Fady, B. (2021). Signature of mid-Pleistocene lineages in the European silver fir (*Abies alba* Mill.) at its geographic distribution margin. Ecology and Evolution, 11(16), 10984–10999.

68. Svenning, J.-C. & Skov, F. (2004). Limited filling of the potential range in European tree species. Ecology Letters, 7, 565–573.

69. Svenning, J.-C., Normand, S., & Kageyama, M. (2008). Glacial refugia of temperate trees in Europe: insights from species distribution modelling. Journal of Ecology, 96, 1117–1127.

70. Stamatakis, A. (2014). RAxML version 8: a tool for phylogenetic analysis and post-analysis of large phylogenies. Bioinformatics, 30(9), 1312–1313.

71. Tinner, W., Colombaroli, D., Heiri, O., Henne, P.D., Steinacher, M., Untenecker, J., Vescovi, E., Allen, J.R.M., Carraro, G., Conedera, M., Joos, F., Lotter, A.F., Luterbacher, J., Samartın, S. & Valsecchi, V. (2013). The past ecology of *Abies* alba provides new perspectives on future responses of silver fir forests to global warming. Ecological Monographs, 83, 419–439.

72. Tzedakis, P. C., Lawson, I. T., Frogley, M. R., Hewitt, G. M., & Preece, R. C. (2002). Buffered tree population changes in a quaternary refugium: evolutionary implications. Science, 297(5589), 2044–2047.

73. Teodosiu, M., Mihai, G., Fussi, B., & Ciocîrlan, E. (2019). Genetic diversity and structure of Silver fir (*Abies alba* Mill.) at the south-eastern limit of its distribution range. Annals of Forest Research, 62(2), 139.

74. Terhürne-Berson, R., Litt, T., & Cheddadi, R. (2004). The spread of *Abies* throughout Europe since the last glacial period: combined macrofossil and pollen data. Vegetation History and Archaeobotany, 13(4), 257–268.

75. Vendramin, G. G., Degen, B., Petit, R. J., Anzidei, M., Madaghiele, A., & Ziegenhagen, B. (1999). High level of variation at *Abies alba* chloroplast microsatellite loci in Europe. Molecular Ecology, 8(7), 1117–1126.

76. Williams, J. W., Shuman, B., Webb, III, T, Bartlein, P. J., & Leduc, P. L. (2004). Late Quaternary vegetation dynamics in North America: scaling from taxa to biomes. Ecological Monographs, 74(2), 309–334.

77. Willis, K. J., & Van Andel, T. H. (2004). Trees or no trees? The environments of central and eastern Europe during the Last Glaciation. Quaternary Science Reviews, 23(23-24), 2369–2387.

